# Prior expectation shapes the emotional response to sounds: behavioural and neural correlates

**DOI:** 10.64898/2026.02.27.708425

**Authors:** Ester Benzaquén, Timothy D. Griffiths, Sukhbinder Kumar

## Abstract

Prior expectations are known to shape perception especially when a stimulus is ambiguous. Bayesian models of cognition posit perception is a precision-weighted combination of top-down and bottom-up information. We consider here affective responses to highly salient stimuli for which a dominant role of bottom-up processing has previously been emphasised. We study how predictions alter the perception of emotional stimuli in a paradigm in which neutral and aversive sounds were preceded by either predictive or non-predictive cues. Cues predicted the type of sound with 100% or 50% probability. Behavioural measures of trial-by-trial expectation and perceived aversiveness were collected before and after stimulus presentation, respectively. We show that prior expectations biased the perceived aversiveness of sounds towards predictions, but only when subjective expectations were considered (as opposed to the objective expectation based on conditional probability). Neural responses were recorded using EEG. During sound processing, we found P3 and LPP components were increased after non-predictive cues, but only for affective stimuli. Time-Frequency results uncovered a role of alpha-beta oscillations in the precision of predictions, as well as in the processing of unexpected stimuli. Our results indicate expectations directly alter the perception of affective stimuli and its processing, and emphasise the importance of behavioural measures to characterize this relationship.

## 1. Introduction

There is now overwhelming evidence that expectation of what is likely to happen in the forthcoming sensory environment shapes perception and neural responses to the stimulus when it arrives (de Lange et al., 2018; Summerfield & de Lange, 2014; Summerfield & Egner, 2009). A consistent finding has been that expectation biases perception towards the expected outcome when the stimulus is noisy and ambiguous. For example, when cued to expect a face, a highly noisy stimulus can be perceived as a face, but when cued to expect a house the same stimulus can be perceived as a house (Summerfield et al., 2006). This can be understood in terms of Bayesian model of perception which posits that perception is a weighted combination of (top-down) expectation and (bottom-up) stimulus features with size of weights determined by the relative precision or reliability of expectation and stimulus. When the stimulus is noisy, its precision is low compared to expectations and therefore the perception tilts in favour of the expected outcome.

Many stimuli in real situations, however, are perceptually noise free, salient and associated with emotional valence (perceived as ‘positive’ or ‘negative’). A large number of studies have shown that stimuli with strong affective valence, when presented unannounced, are prioritized (compared to neutral stimuli) while processing both at the behavioural (Ohman et al., 2001; Phelps et al., 2006) and neuronal level (Anderson et al., 2003; Pessoa et al., 2002). All these studies point to a dominant role of bottom-up processing of affective stimuli in that the features of stimuli alone determine behavioural and neural responses. Many a time, however, the onset of an affective stimulus is predicted by a cue or the context in which the stimulus is presented. For example, seeing somebody standing near a blackboard with a chalk in his/her hand, expectation of a screeching sound can be generated by this visual cue. How does this top-down factor of expectation influence the perceived and brain response to the affective stimulus?

Expectation influences both pre-stimulus and post-stimulus behavioural responses. For example, a feeling of dread is experienced during the time interval that precedes the onset of a negative stimulus (Berns et al., 2006). Similarly expectation of a positive event initiates positive feelings during the delay phase (Golub et al., 2009). How the expectation weighs-in on the affective response post-stimulus onset is less clear. A large body of evidence from pain perception (Atlas & Wager, 2012) shows that the response to a stimulus is shaped towards the expectation: that is, a mild stimulus is perceived as more painful if the subject expects a highly painful stimulus compared to when the same stimulus is presented following expectation of a mild or moderate pain. Other studies show that expectation either biases the response towards it or has no effect on the affective response following a stimulus (for a review, see Golub et al., 2009). Since the affective stimulus is highly salient and unambiguous, the high (bottom-up) precision accorded to the stimulus competes with the precision of predictions. That is, weighting of priors in perception can be more, less or equal to the weighting of stimuli. This has implications for studies investigating the neural basis of expectation in that there could be variation in the effect across subjects: some participants can hold on to their expectations with high precision and their percept would be biased toward priors whereas others may not be affected by the expectation.

A typical paradigm to investigate the role of expectation on affective processing involves presenting affective stimuli following a cue with perfect certainty (100% cue validity) while in another condition affective stimuli are presented after a non-predictive cue (50% cue validity). One problem with these type of design is that objective (conditional) probabilities of the affective stimulus following a cue (100% in certain and 50% in uncertain) are assumed to be the same as the subjective certainty experienced by participants: just because a cue predicts the stimulus with 100% validity does not mean that participants ‘feel’ certain that stimulus will occur. In addition, when the cue is uncertain, it does not mean that people do not make a prediction about the upcoming stimulus (Catena et al., 2012). As explained previously, this distinction is particularly important for affective salient stimuli because of the close competition between expectation and stimulus salience. To separate the objective and subjective expectation, it is important to measure individual subject’s expectations before stimulus onset.

Theoretical and empirical research under the Bayesian framework argues that predictions associated with highly precise priors reduce neural activity during stimulus processing (Friston, 2005). This is clearly illustrated by the results of neural adaptation or repetition suppression (Desimone, 1996). Research on human affect has prominently involved two distinct but related ERP components, the P3 or P300, and the late positive potential (LPP), both reflecting the cognitive saliency of emotional or motivational stimuli (Hajcak et al., 2010; Hajcak et al., 2012). Prediction effects on ERPs in response to emotional stimuli have yielded inconsistent results. For example, Lin et al. (2015) found an enhanced P3 for unpredicted positive images and decreased LPP for unpredicted aversive images. Others have found enhanced LPP for non-predicted conditions during processing of both neutral and negative pictures (Dieterich et al., 2016; Johnen & Harrison, 2019). Most studies fail to record subjective expectations, while participants are explicitly informed of the objective probabilities, which may alter how these predictions are formed and internalised.

Top-down predictions under the Bayesian framework are considered to be represented by oscillatory activity in the beta-frequency band (Bastos et al., 2012; Sedley et al., 2016). However, decreases in alpha oscillations have also been associated with temporal predictions (Rohenkohl & Nobre, 2011; Thut et al., 2006), and alpha suppression is proposed to reflect top-down control from attentional networks (Mathewson et al., 2014). Affective research has shown that activity in the lower frequencies, such as theta, alpha and beta frequency bands, can code for top-down modulation and expectations of pain (Babiloni et al., 2003; Huneke et al., 2013; Strube et al., 2021b; Taesler & Rose, 2016; Tiemann et al., 2015; Tu et al., 2016), as well as aversive images (Strube et al., 2021a). Whether activity in alpha-beta or lower frequency bands embodies predictions, the precision of predictions, or both, remains unexplored.

In the current study we measured behavioural and neural responses while subjects expected the onset of aversive or neutral sounds following certain or uncertain cues. We used scratchy sounds (e.g. sound of knife scratched on bottle) which have been characterized acoustically (Kumar et al., 2008) and are perceived as highly aversive and salient by most. The role of expectations on perceived aversiveness, as well as on the neural processing of aversive sounds and their prediction are examined.

## 2. Materials and Methods

### 2.1. Participants

Twenty-five participants were recruited to take part in this study (15 females; mean ± SD: 25.30 ± 5.79 years; age range: 18 – 37). All participants were right-handed, had normal hearing and normal or corrected-to-normal vision, and no history of neurological or psychiatric disorders. The study was approved by Newcastle University Research Ethics Committee (Reference number: 1418/732/2017). Written informed consent was obtained from all participants prior to the start of the experimental protocol. All volunteers were provided with an inconvenience allowance of £15 for their participation.

### 2.2. Stimuli

The stimuli consisted of a set of ten one-second sounds of two categories: aversive and neutral. All five neutral sounds were water-sounds chosen for their lack of unpleasantness and inability to induce feelings of threat or anticipatory fear. All unpleasant sounds were scraping or scratching sounds from previous work developed by Kumar et al. (2008). Two of the neutral sounds were from that same source (Kumar et al., 2008), while one was from the International Affective Digitized Sounds (IADS-2; Bradley & Lang, 2007). The remaining two sounds were collected from a free internet source (https://www.zapsplat.com/). Thus, these two neutral sounds were not acoustically or behaviourally characterised. The open-source software Audacity® (https://www.audacityteam.org/) was used to edit stimuli duration. When necessary and using the same software, sounds were further altered to resemble each other in loudness and sampling rate.

### 2.3. Experimental protocol

The task structure followed that of a trace conditioning paradigm. Participants were presented three different visual cues: **‘X’**, **‘Y’**, and **‘?’**. In all cases, ‘X’ was followed by an aversive sound, whereas ‘Y’ was followed by a neutral sound with 100% cue validity. In the third condition, the visual cue **‘?’** was followed by either a neutral or aversive sound, such that half of the trials consisted of aversive sounds and the other half of neutral sounds. Hence, sounds could be accurately predicted by the cue (first and second conditions; ‘certain trials’) or not (third condition; ‘uncertain trials’). Participants were not made aware of these contingencies but were told to use the cues to try to predict which type of sound they were going to hear.

After the presentation of the cue, participants reported the likelihood that an aversive sound would follow on a scale from 0 (not at all l, i.e. a neutral sound was expected) to 100 (most certainly). Participants were instructed to move a slider using the computer mouse and to respond within a fixed period of two seconds. An anticipatory interval of four seconds was introduced after the likelihood rating which was followed by a sound of 1s duration. After sound offset, participants rated the aversiveness of the preceding sound on a scale from 0 (not at all) to 10 (most aversive) within a maximum of 3 seconds. A new trial started 3 seconds after a response was made (ITI; inter-trial-interval). The trial structure and timeline are depicted in Figure 1.

**Figure 1.**
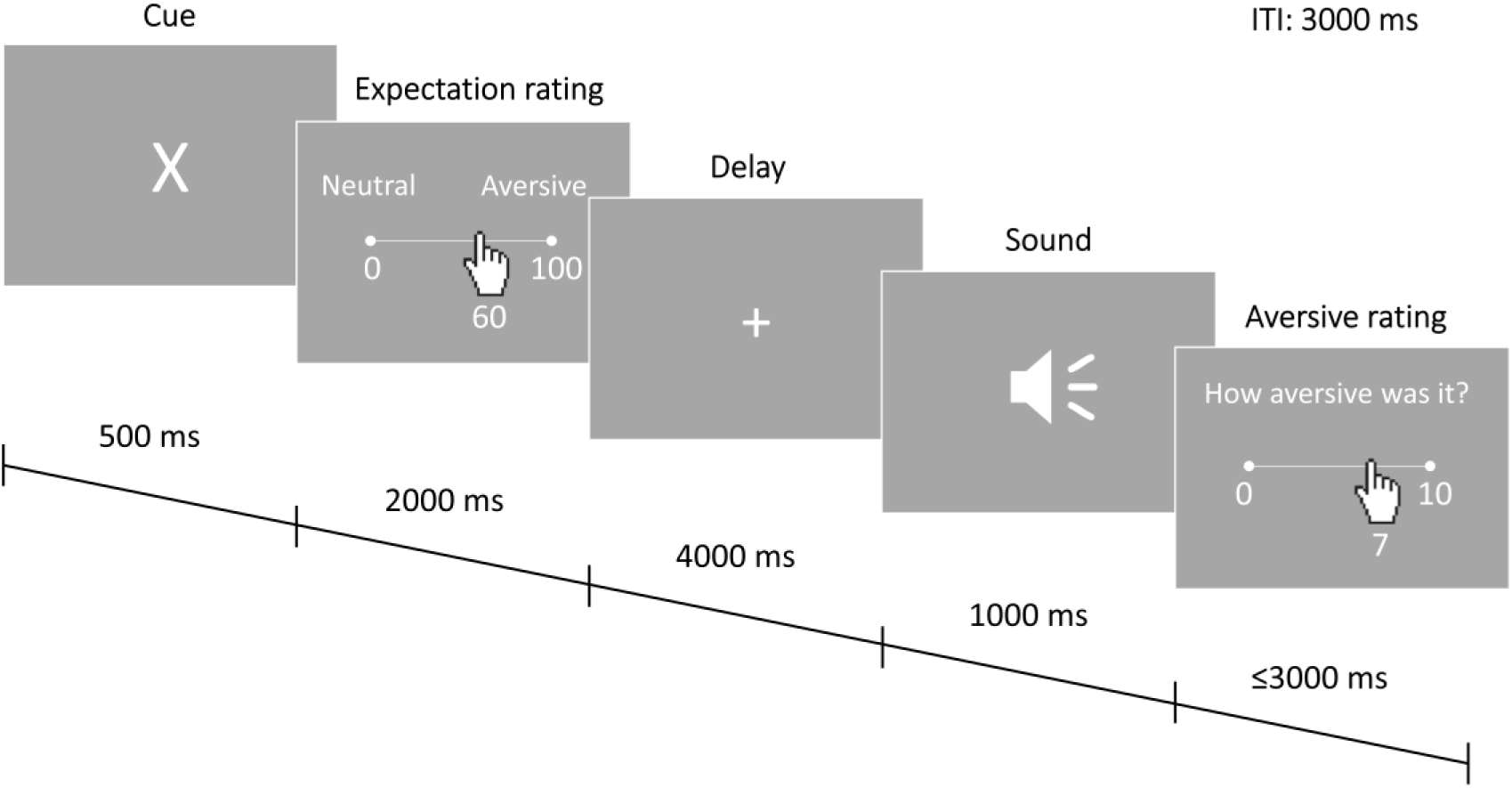
Cues were displayed for 500 ms, followed by an online rating on how likely participants thought that the subsequent sound would be aversive. Participants had to answer by moving a slider between 0 and 100 during a 2 s period after the offset of the cue. A delay of 4 s where a fixation cross was displayed was immediately followed by the onset of an either aversive or neutral sound. After stimulus offset, participants rated the aversiveness of the stimulus within 3 seconds on a 0-10 scale. The trial was terminated as soon as participants responded, and another trial started 3 seconds later (Inter trial interval [ITI]).

The task, with 80 trials per condition presented in a pseudorandomised order, was divided into 4 blocks, with rest periods offered between them. Sound presentation was fully randomised within conditions, and each sound was played 24 times. Before the experimental sessions, participants completed a brief practice session where all three cues were presented twice in a fixed order. Two sounds different from those used for the experimental task but with similar characteristics were used (i.e., one water- and one scraping-sound) during the practice session. The entire study session, including set-up time, lasted approximately two hours.

### 2.4. EEG measurements

EEG was recorded using a 64-channel ActiveTwo BioSemi system (BioSemi Instruments, Amsterdam, The Netherlands). Vertical and horizontal EOG (electro-oculogram) signals were recorded from electrodes positioned above and below the right eye and on the outer canthi of both eyes. Electrode DC offset was kept stable and within ±30 mV.

### 2.5. Behavioural data analysis

Expectancy ratings and their reaction times (RTs) were averaged for each condition, and then compared across conditions (aversive, neutral, uncertain) using repeated measures ANOVA (analysis of variance). Reaction times for the expectancy ratings were missing for 2 participants due to a technical issue. Sphericity was assessed with Mauchly’s test, and Greenhouse-Geisser correction was applied when this assumption was violated (*p* < .05). Aversive ratings were first standardized (mean = 0, standard deviation = 1) across all trials and the average value for each condition was calculated for each participant. A 2×2 repeated measures ANOVA with factors ‘valence’ (unpleasant and neutral) and ‘certainty’ (predictable and unpredictable) on expectancy ratings and reaction times was performed to assess the effect of certainty and valence and their interaction.

To determine whether a change in valence ratings is mediated by individual’s reported expectation, mediation analysis (Baron & Kenny, 1986) was performed using the Mediation Toolbox (https://github.com/canlab/MediationToolbox) on non-standardised ratings. Briefly, mediation analysis assesses if the relationship between an experimental manipulation (X), and the outcome or dependent variable (Y), can be explained by a third variable (M). In our case, the independent variable ‘X’ consisted of the certainty or cue manipulation [certain = 1, uncertain = -1], the outcome ‘Y’ were the reported aversive ratings, while ‘M’ were the reported expectancy ratings. The model defines different paths (*a*, *b*, *c*, *c’*, *ab*) from ‘X’ to ‘Y’ to test for the relationship between the three variables using a series of ordinary least squares regressions. In our case, path *a* (X -> M) represents the certainty effects on expectation. The effects of expectation on aversive ratings (M -> Y), path *b*, is measured while controlling for X. Path *c* and *c’* represent the effect of X on Y, before and after controlling for the effect of M, respectively. The combined path, *ab*, defined as the product of *a* and *b*, tests whether the introduction of M accounts for the relation between X and Y. Statistical significance was established with a bootstrap test using 10000 permutations as performed in the Mediation Toolbox. The mediation analysis was performed separately for aversive and neutral sounds.

### 2.6. EEG data reduction and analysis

#### 2.6.1. Preprocessing

Data preprocessing was performed using EEGLAB software (https://www.sccn.ucsd.edu/eeglab/), while statistical analysis was accomplished with FieldTrip (http://fieldtriptoolbox.org/). First, data were low-pass filtered to 100 Hz and downsampled from 1024 to 256 Hz. Trials were epoched and detrended (de Cheveigne & Arzounian, 2018) to remove slow drifts instead of the standard application of a highpass filter, as filtering may introduce distortions in the data (Tanner et al., 2015; Yael et al., 2018). Channels poorly correlated (r < 0.8) with their RANSAC (random sample consensus) reconstruction were rejected and interpolated (mean ± SD: 5.64 ± 3.74 channels). Ocular-artifacts were identified using AMICA (Adaptive Mixture Independent Component Analysis) (Palmer et al., 2011), which achieves a better ICA decomposition compared to other algorithms such as Infomax (Delorme et al., 2012). Horizontal and vertical bipolar EOG signals were calculated by subtracting each horizontal and vertical recorded channel together, respectively. A total of 32 components were extracted and correlated with bipolar EOG signals. A component was considered an ocular artifact when this correlation exceeded 0.6 and confirmed by further visual inspection. Data were referenced to common average and entire trials were epoched from -3 to 10.5 seconds from cue onset. Baseline correction was performed 100 ms before cue onset [ERP analyses only] and trials were further epoched into different conditions with different timings depending on analysis. For sound ERP analysis, epochs were formed from 200 ms before sound onset to 200 ms after sound offset [-0.2 1.2] and divided into 4 conditions depending on cue and sound type (aversive certain, aversive uncertain, neutral certain, neutral uncertain). For time-frequency analyses, trials were epoched from 3 seconds before cue onset to sound offset and divided into 3 conditions depending on cue (aversive, neutral, uncertain). Further artifacts were rejected from analysis by identifying trials outside 5 SDs from the mean and whose kurtosis was larger than eight. To prevent the introduction of rejection bias towards less common trials (i.e., uncertain), trial rejection was performed after epoching each condition and for each analysis separately. This resulted in the rejection of (mean ± SD) 6.24 ± 3.7% of all trials across all subjects.

#### 2.6.2. ERPs

ERPs were baseline corrected from 100 ms before cue onset and low-pass filtered to 30 Hz. Statistical analyses were performed using a Monte Carlo permutation test with 1000 permutations and a cluster correction with a threshold of *p* < .05.

Sound ERPs were baseline corrected using 100 ms pre-sound onset. The P3 component was visually identified on sound evoked ERPs. To avoid bias (Luck, 2005), ERPs were averaged across all conditions and the quadratic mean [RMS; root mean square] over all channels was calculated. The P3 component, which peaks around 300 ms, was then visually identified in the created RMS wave. Statistical analyses were performed over the entirety of the selected P3 time-window [246 – 348 ms] and over all channels. The LPP was divided into 200 ms-long windows, starting at 400 ms after sound onset, resulting in three time-windows [400 – 600 ms; 600 – 800 ms; 800 – 1000 ms]. Statistical analyses were performed for each time-window separately.

#### 2.6.3. Time-Frequency analysis

Using EEGLAB, time-frequency (TF) decomposition was performed with complex Morlet wavelets. TF activity from 2 to 47 Hz was calculated using linearly increasing cycles: starting at 3 cycles at the lowest frequency and increasing with higher frequencies to reach 8 cycles. Time-frequency data were baseline corrected using the time period from 1.2 to 0.2 seconds before cue onset, and TFRs (time-frequency responses) were created during cue presentation by averaging TF power [2-47 Hz] over all channels and time (0-0.5 seconds timelocked to cue onset).

The role of expectations on TF power was explored by comparing TFRs during certain (i.e., aversive and neutral) and uncertain cues, again using Monte Carlo permutations (n=1000) and cluster-correcting for multiple comparisons.

The relation between time-frequency power and expectations was assessed using a data-driven approach. First, expectancy ratings were transformed to reflect the precision or confidence of predictions. An uncertainty index (UI) was created to compute the distance to the extreme ends of the expectancy scale, such that greater numbers signify less confident predictions (closer to the middle of the scale). This was calculated as the absolute difference between the reported expectancy and the real probability, such that UI = | *Probability_REPORTED_* – *Probability_REAL_* |. So, for example, if a participant was confident that aversive sounds would play after the aversive cue, and they rated this as ‘99’, the uncertainty index (UI) would be ‘1’, as |99 - 100| = 1. If, on the other hand, they were less confident that a neutral sound would follow the neutral cue, and their averaged expectancy rating was ‘20’, the UI would be ‘20’, as |20 - 0| = 0. This allows for a direct comparison between expectations during neutral and aversive trials, since a completely certain expectation during aversive trials (aversive expectation = 100) and during neutral trials (aversive expectation = 0) would have equal uncertainty indexes (UI = 0 in both cases).

To test which frequency bands correlated with expectation, time- and channel-averaged TFRs during cue presentation were correlated with the UIs using Pearson and cluster-corrected for multiple comparisons using 1000 Monte Carlo permutations. To further explore the role of certainty on TF power, the same method was followed using the standard deviation of said expectancy ratings. Greater SDs in expectancy rating were considered as less precise predictions, and thus imply greater uncertainty. To assess whether established correlations over significant frequencies during cue presentation and transformed expectancy ratings (i.e., UI) were a corollary to the upcoming motor response of such expectancy ratings and thus represent motor preparedness, the same correlation method was repeated but with individual’s mean RTs.

## 3. Results

### 3.1. Participants

Data were collected from 25 participants. Participants who failed to learn the cue-sound association were excluded from all analyses to remove any spurious effects they could cause. This was defined as a mean aversive expectation ≤ 60 for the aversive cue or ≥ 40 for the neutral cue. A sample of 22 (12 females; age: 24.79 ± 5.42 (mean ± SD); range 18-37) participants remained. There was no difference in age (*t*(45)=0.32, *p*=0.75) nor sex (χ^2^(1) = 0.14, *p* = 0.71) between the original and final samples.

### 3.2. Behavioural results

#### 3.2.1. Expectancy

Remaining participants learnt the contingencies as shown by their averaged expectancy responses (mean ± SD: aversive = 88.9 ± 11.2; neutral = 12.5 ± 13.3; uncertain = 52.9 ± 8.8; Figure 2A), which differed between conditions (F(1.32,42) = 193.2, *p* < .001, η^2^ = .902; Greenhouse-Geisser corrected). Pairwise comparisons demonstrated that all three conditions differed (*p* < .001) from each other. Expectancy reaction times were also different between conditions (F(1.238,20) = 34.24, *p* = .000, η^2^ = .643; Greenhouse-Geisser corrected; Figure 2B). Pairwise comparisons showed that participants responded over 200 ms slower during the uncertain condition compared to both certain conditions, demonstrating induced uncertainty (mean ± SD: uncertain = 1241 ± 121 ms; aversive = 990 ± 167 ms [*p* < .001]; neutral = 1018 ± 160 ms [*p* < .001]).

**Figure 2.**
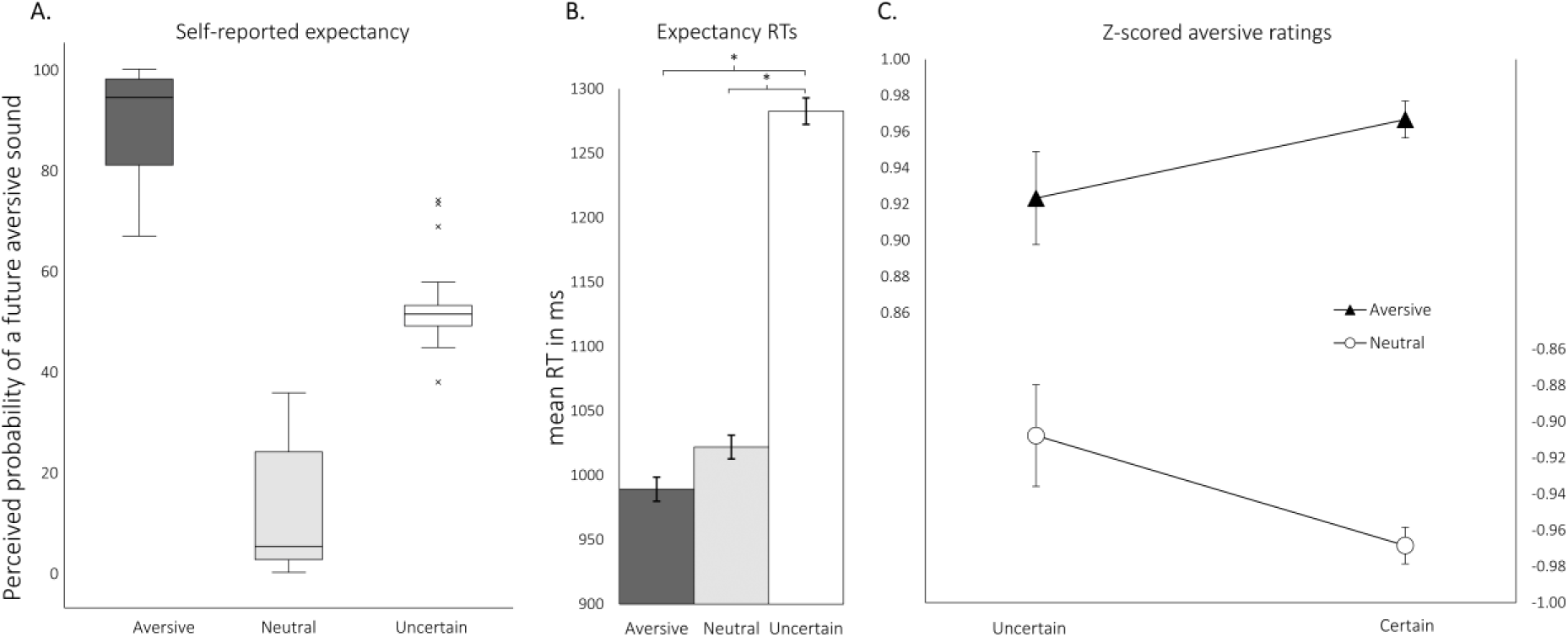
A. Self-reported expectancy (probability of an aversive sound) after aversive, neutral, and uncertain cues. B. Mean reaction times of expectancy ratings. C. Mean standardized aversive ratings for aversive (left Y-axis) and neutral (right Y-axis) sounds following certain and uncertain cues. Greater values imply greater aversiveness. Bars represent SEM (standard error of the mean). * = p < .001.

#### 3.2.2. Aversive ratings

A repeated measures two-way ANOVA examined the effects of certainty and valence on aversive ratings, which showed main effects of valence (F(1,21) = 6705, *p <* .001, η^2^ = .997) but no main effects of certainty (F(1,21) = 0.94, *p* > 0.05, η^2^ = .043). The interaction was not significant (F(1,21) = 2.64, *p* = 0.1, η^2^ = .112) implying that expectation did not influence the perception of aversive and neutral sounds differently (aversive sounds becoming more aversive and neutral sounds becoming less aversive when expected). A trend, however, in this direction was observed (aversive: certain= .97 ± .05, uncertain = .92 ± .1; neutral: certain = - .97 ± .05, uncertain = -.91 ± .13; Figure 2C).

#### 3.2.3. Relation between expectancy and aversive ratings

To explore the role of individual’s predictions in the aversive experience, we performed a mediation analysis with participant’s predictions (expectancy ratings) as a mediator variable. The results (see Figure 3) revealed that expectancy mediated the shift in perception significantly (*p* < 0.001) for both aversive and neutral sounds. As shown in the scatter plots of Figure 3, expectation ratings positively correlated with aversive ratings, such that greater expectancy for an aversive sound resulted in greater reported aversiveness. Similar results were obtained for the neutral sounds, such that greater expectancy of a neutral sound resulted in less reported aversiveness.

**Figure 3.**
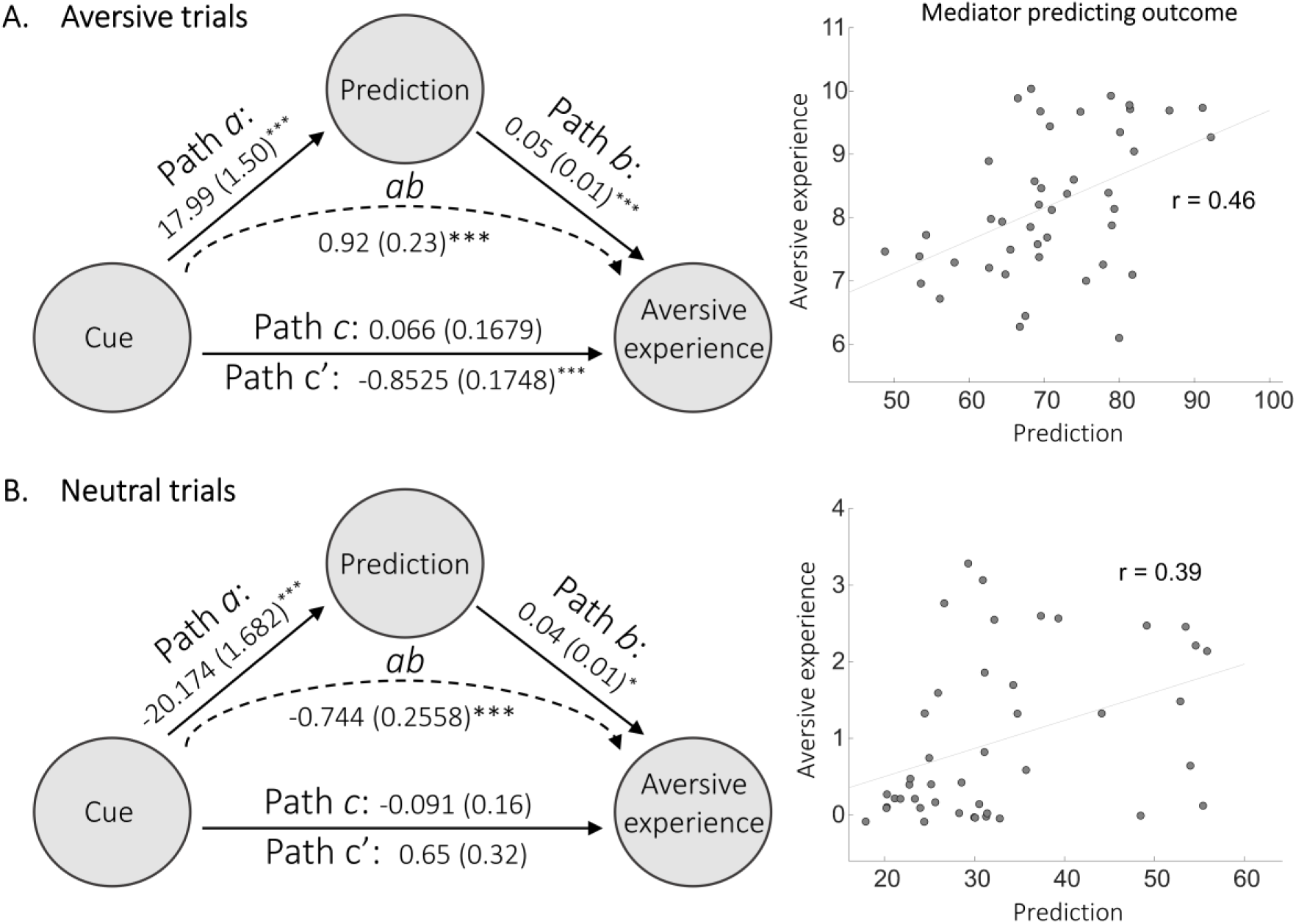
Mediation analysis results between cue type (1: certain; -1: uncertain), prediction, and reported aversive experience. Left panels show mean path coefficients with standard errors in parentheses for aversive (A) and neutral (B) trials. *** = p < .001; * = p < .05. Right panels show partial regression scatterplots for the relation between mediator (prediction) and outcome (aversive experience) controlling for cue effects.

### 3.3. ERP results

#### 3.3.1. Sound period

Aversive sounds that could not be accurately predicted, that is, those preceded by the uncertain cue, elicited a greater P3 wave over central electrodes throughout the analysed time-window [246 – 348 ms] (Figure 4A). This difference, although still present, did not achieve statistical significance during neutral sounds. Similarly, the early-LPP [∼500 – 600 ms] exhibited an increased amplitude during uncertain sounds compared to certain sounds, but only aversive trials achieved statistical significance (Figure 4B). This difference was absent in later stages of the LPP. Scalp topographies of the difference between aversive sounds (uncertain – certain) and ERPs of selected channels are depicted in Figure 5.

**Figure 4.**
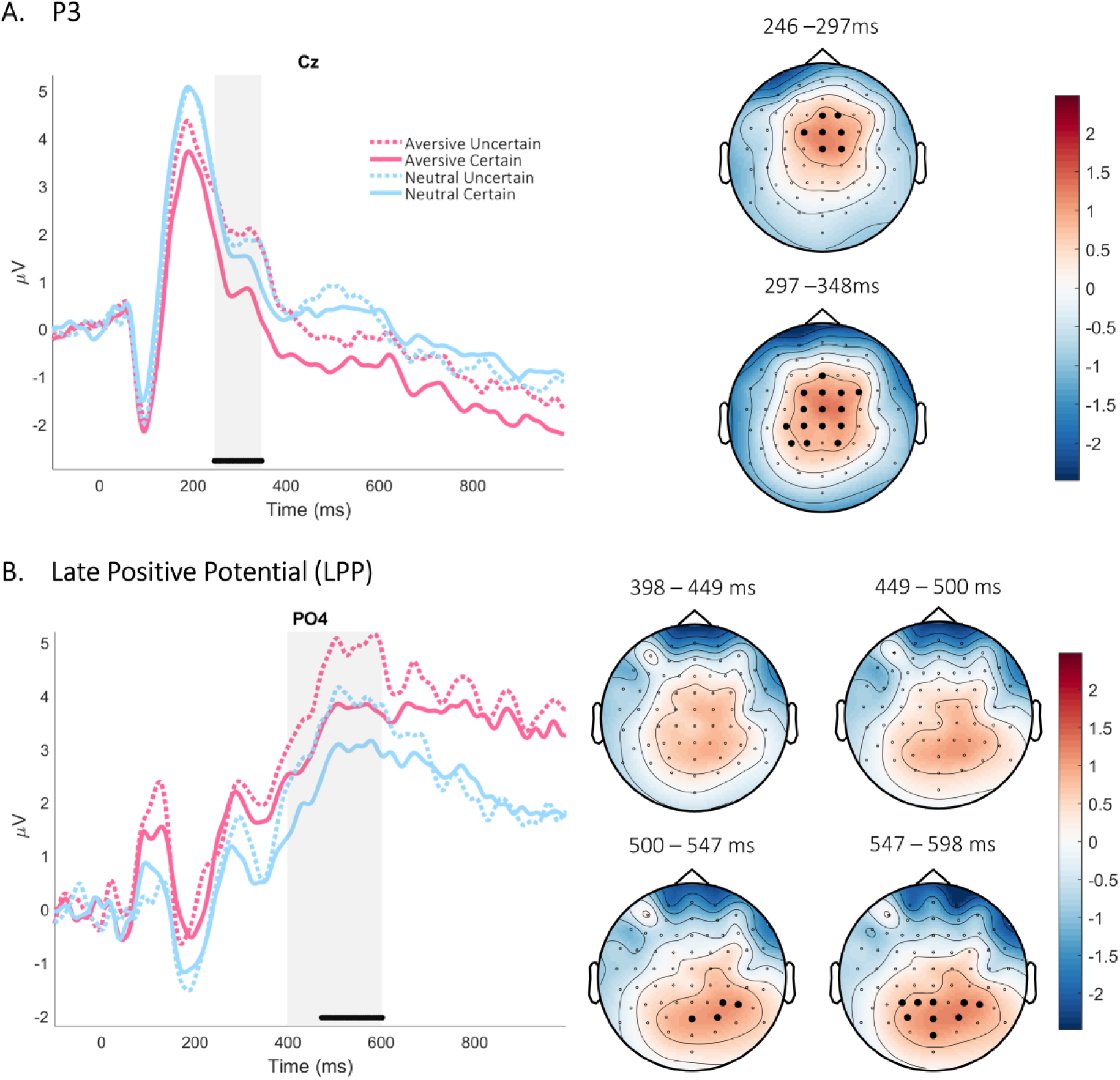
Sound evoked potentials. Difference maps between uncertain and certain aversive sounds on the right, and ERPs on selected channels on the left. The P3 time-window [246 – 348 ms] was identified and analysed (A). The LPP was divided in three 200 ms-long windows [400 – 600; 600 – 800; 800 – 1000 ms]. Significant differences were found on the earlier time-window (B). Grey shaded areas represent analysed time windows. Black dots indicate significance between aversive certain and uncertain sounds (p < .05; cluster corrected). No significant differences were found between certain and uncertain neutral sounds.

**Figure 5.**
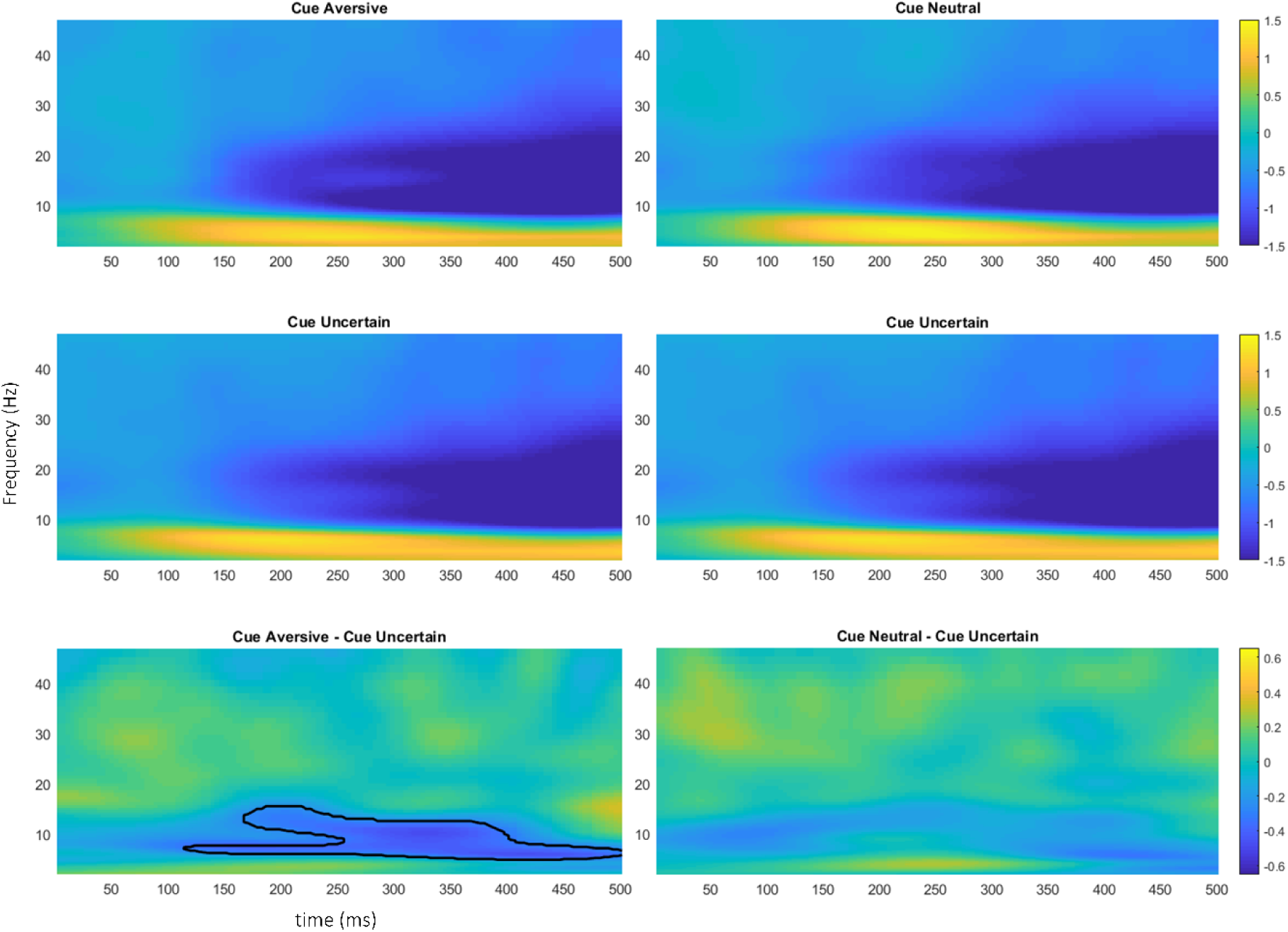
Time-frequency results during the cue period. Lower panel shows the difference in power between certain (aversive: left, neutral: right) and uncertain cues. Significant clusters (p < 0.05) are highlighted in black.

### 3.4. TF results

#### 3.4.1. Cues

Cue presentation was characterized by an increase on lower frequencies (theta: θ), and alpha (α) suppression that extended to the high beta-band (β). When comparing aversive and uncertain cues, a significant negative cluster between 5-16 Hz was found, meaning power over these frequencies was greater on the uncertain condition. Thus, θ power increased after uncertain cues, and α suppression was enhanced after certain aversive cues. These results can be seen in Figure 5.

#### 3.4.2. Relation between TF power during cues and expectancy ratings

Significant clusters (*p* < 0.05) were found when correlating averaged power during the presentation of certain cues (i.e., aversive and neutral cues) and transformed expectancy ratings (uncertainty index: UI), as can be seen in Figure 6A. To better characterise these results and for illustration purposes only, power over cluster-identified frequencies was collapsed (averaged) and correlated with UIs, which yielded correlations over the beta-frequency band. Greater certainty of an aversive sound after an aversive cue was coupled with reduced beta power (17-34 Hz; r = 0.553; Figure 6B). Similarly, enhanced confidence that a neutral sound would play after the neutral cue was also paired with reduced beta power (15-22 Hz; r = 0.569; Figure 6B). Thus, the degree of certainty of predictions translated into an increase in beta suppression. To further ascertain the role of certainty on induced oscillations, frequency power over all frequencies was again correlated with the standard deviation of expectancy ratings (Figure 6C). Significant clusters were found over the alpha- and beta-band. Again, power over clustered-frequencies was averaged and repeatedly correlated with SDs, which produced a positive correlation for aversive (12-25 Hz; r = 0.585) and neutral cues (9-26 Hz; r = 0.652; Figure 6D). Thus, less uncertainty, and hence less deviation in responses, was coupled with greater alpha-beta suppression. To reject the possibility that this effect was driven by a preparatory motor response due to the fact that expectation ratings were gathered immediately after cue offset, power during cue processing was correlated with participant’s reaction times using the same approach. This yielded no significant result. For comparison, power over frequencies that were part of significant clusters in the correlation with UIs were averaged and correlated with RTs, which again showed no significant correlation for either aversive (17-34 Hz; r= 0.254, *p* > 0.2) or neutral (15-22 Hz; r= 0.235, *p* > 0.3) trials. No significant correlations were found between TF data during presentation of the uncertain cue and expectancy ratings, their SDs, or RTs.

**Figure 6.**
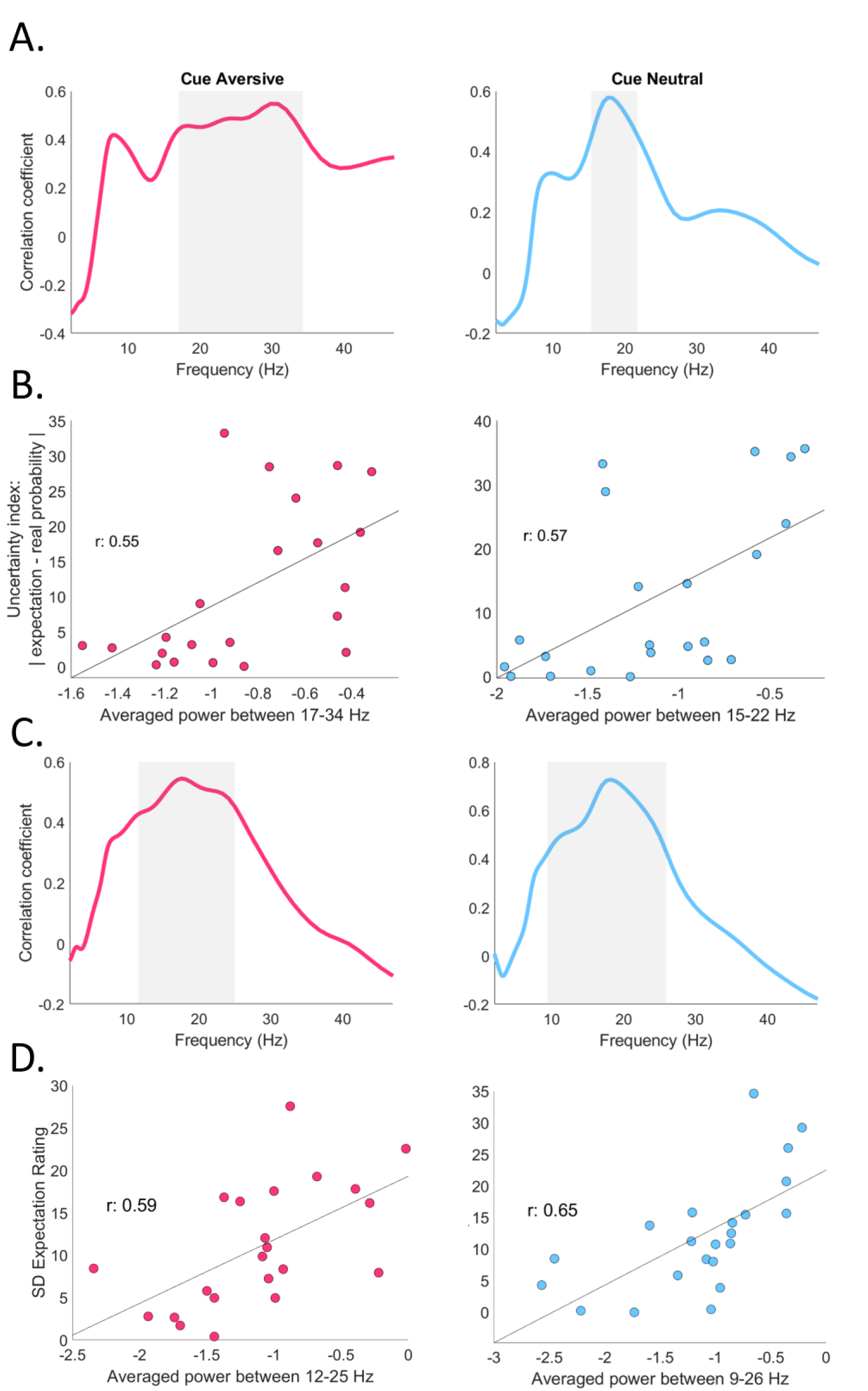
Correlations between TF power during cue processing and subjective expectation. A. TF power was correlated with an uncertainty index (UI) representing the strength or confidence of predictions. Significant frequency clusters (p < 0.05, cluster corrected) are highlighted in grey. B. Correlations between TF power over significant frequencies and UIs (for illustration). C. TF power was correlated with the SD of expectancy ratings and significant frequency-bands highlighted in grey (p < 0.05, cluster corrected). D. For illustration, power over significant frequencies was correlated with SDs.

#### 3.4.3. Sounds

Sound onset was characterised by an increase on delta-theta power (∼2-7 Hz) that progressively dissipated. This delta-theta power was greater during neutral sounds that were preceded by the uncertain cue compared to neutral sounds accurately predicted by a neutral cue. No differences between aversive sounds according to preceding cue were found on delta-theta power. Around 400 ms, an alpha-beta (∼6-24 Hz) power suppression was observed, which was stronger for aversive sounds that were not predicted by the cue. This difference, although present during neutral sounds, did not reach statistical significance. These results can be seen in Figure 7.

**Figure 7.**
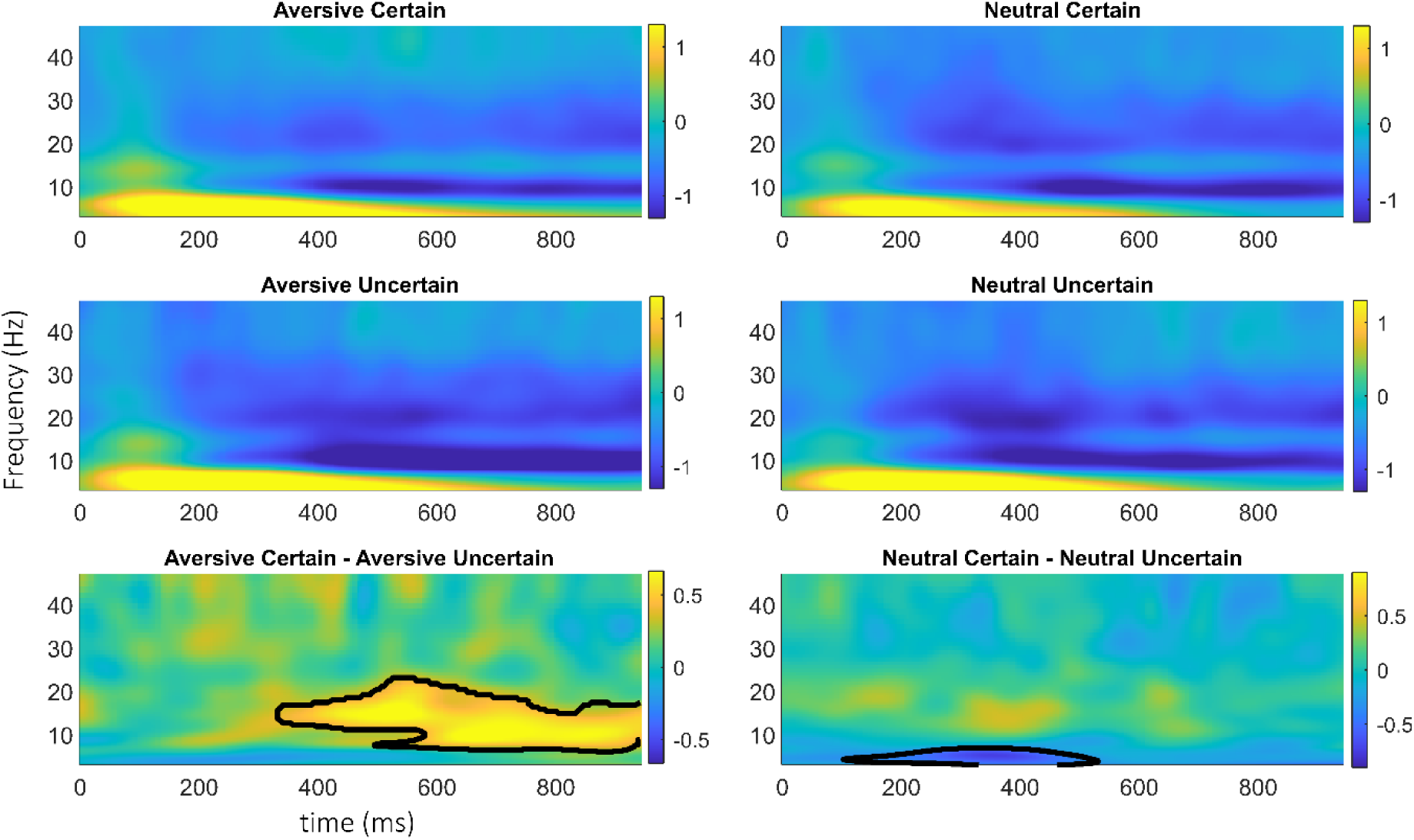
TF results after sound onset. Lower panel shows the difference in power between certain and uncertain sounds. Significant clusters (p < 0.05, cluster corrected) are highlighted in black.

Power over the β-band was further explored by averaging over said frequency-band (13-30 Hz). Aversive and neutral sounds were compared according to the preceding cue, which revealed that β-suppression was stronger for uncertain sounds at around 400 ms, as can be seen in Figure 8. This difference, although short-lived during neutral sounds, extended beyond 700 ms during aversive sounds. Similar exploration of the θ-frequency band (3-8 Hz) uncovered a dissociation according to valence: aversive uncertain sounds showed θ-decrease when compared to aversive certain sounds, while uncertain neutral sounds displayed enhanced θ-power when compared to certain neutral sounds. Further, this latter difference emerged at ∼200 ms, while the difference in aversive sounds only occurred at around 800 ms post-sound onset (see Figure 8).

**Figure 8.**
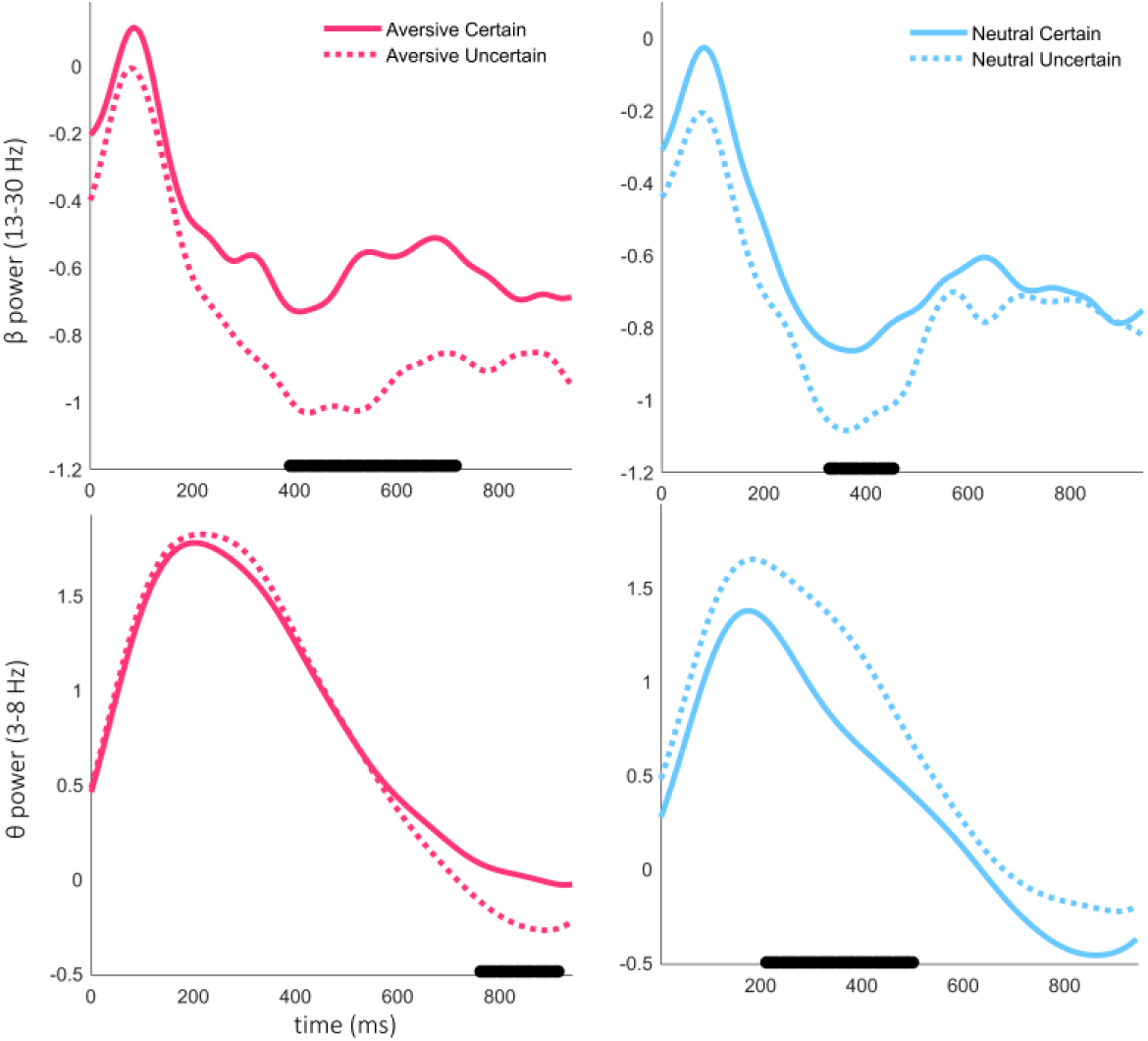
TF results over selected frequency bands during sound processing. Significant time clusters are highlighted in black (p < 0.05, cluster corrected).

## 4. Discussion

Here, we manipulated the predictability of aversive and neutral sounds while recording measures of subjective prior expectancy and sound aversiveness as well as brain measures using EEG. Two predictive cues preceded aversive or neutral sounds, respectively, while a third non-predictive or uncertain cue preceded both aversive and neutral sounds in equal proportions. Our results show that even though the predictability manipulation did not significantly alter participants’ reported aversiveness to sounds at the group level, taking into consideration individuals’ predictions revealed expectations played a mediator role between the cue and the perceived aversiveness. This indicates that the experimental manipulation of expectation affected participants on an individual basis. In terms of Bayesian theory of perception, it means that precision of prior expectations following the same cue varied across subjects. This individual variation in precision may not be discernible when the upcoming stimulus is ambiguous or noisy but is strong enough to mask the expectancy effect at the group level when the stimulus is salient. Taking into consideration an individual subject’s expectation using explicit behavioural measures can reveal expectancy bias on the percept. This is in line with previous research in pain processing, which showed pain ratings were biased towards individuals’ priors (Zaman et al., 2017), as well as similar work to the current study performed in the visual domain (Lin et al., 2012; Lin et al., 2015). Our study adds to a growing body of literature showing different aspects of sensory cognition are biased in line with expectations, from visual perception (Aitken et al., 2020) to action outcomes (Yon et al., 2020), and even higher-level affective processing as shown here.

EEG measures showed neural responses differed between sounds preceded by certain and uncertain cues. We focused here on components P3 and LPP due to their involvement in affective processing (Hajcak et al., 2010; Hajcak et al., 2012). Previous work show that P3 and LPP are affected by the saliency or arousal (Dillon et al., 2006; Schupp et al., 2000), self-relevance (Gray et al., 2004), and emotional content (Johnston et al., 1986) of the stimuli. Here, we demonstrate that the same aversive sounds elicit a greater P3 and LPP when preceded by an uncertain non-predictive cue. Since bottom-up information remains the same, this difference in auditory ERPs can only be due to top-down effects of expectation. Most importantly, top-down expectation did not alter the same components for neutral sounds. The apparent expectation effect only during the processing of aversive sounds may imply involvement of brain structures such as insula and amygdala specifically for aversive stimuli which is also consistent with fMRI research (Sarinopoulos et al., 2010) and the finding that generators of LPP are found to be in insula particularly for negative affect stimuli (Liu et al., 2012). P3 is commonly studied with oddball paradigms, where uncommon probes or stimuli elicit a greater positive deflection around 300 ms compared to frequent stimuli. The P3 is thus considered to signal surprise or novelty and represents shifts in attention, and insula has been implicated as one of its sources (Horovitz et al., 2002). The involvement of the insula on the processing of unpredictable affective stimuli is further supported by fMRI research (Ran et al., 2018). We are, unfortunately, unable to separate expectation and attentional effects, and it remains a possibility that some of our results are confounded by increased attentional capture by affective stimuli.

Data-driven analyses demonstrated the specific role of alpha and beta oscillations in top-down effects of expectations. During cue processing, an alpha-beta suppression was observed, which was enhanced for the processing of cues with predictive value (certain). Further analyses showed oscillatory activity in the alpha-beta band encoded the strength or precision of predictions, such that greater certainty (precision) was associated with greater alpha-beta suppression during cue-encoding, supporting previous evidence relating alpha (Sedley et al., 2016) and beta (Palmer et al., 2019) oscillations with prediction precision. Oscillatory activity in the beta frequency range is modulated by movement (e.g., Pfurtscheller & Lopes da Silva, 1999); thus, to reject the possibility that recorded beta-band activity during cue processing reflected upcoming motor responses, correlations between TF power and reaction times were performed, which revealed no associations.

Effects of predictions on sound processing were again indexed by oscillatory activity in the beta band. Aversive and neutral sounds not predicted by the cue elicited greater reductions in beta power. This and the previous results indicate that beta-power suppression appears to index the precision of predictions and what could be considered a ‘prediction error’. These apparent opposite results can be unified if beta-power desynchronization is considered to also reflect an updating of predictions (Sporn et al., 2020), or simply the encoding of stimuli (Lundqvist et al., 2016). It can also be argued that activity in the beta frequency band might correspond to different processes, maybe depending on source (Schmidt et al., 2019), which was not explored here. However, a seemingly more parsimonious explanation could be that alpha-beta suppression signals control of attentional resources, as has been proposed by others (Klimesch, 2012; Wiesman & Wilson, 2020). In this case, participants who are better at recruiting attentional processes (indexed by reductions in alpha-beta power) will learn cue-sound contingencies more accurately, thus showing greater ‘precision’ of predictions.

It is worth noting, however, that the use of a non-predictive cue that was not devoid of meaning, in this case a question mark (‘?’), may be behind some of the previously discussed results. Unlike the predictive cues (‘X’ and ‘Y’), ‘?’ did not necessarily require associative learning, as it was likely considered to signal an unknown outcome due to its semantic meaning. Therefore, this difference between predictive and non-predictive cues is a major limitation of the current study.

To summarise, the current study demonstrates that expectations can modify the subjective experience to affective sounds in the direction of predictions showing an expectancy bias in perception, but only when individual predictions were considered. Further, unpredictability enhanced (or predictability attenuated) components P3 and LPP only during exposure to aversive sounds. Time-frequency analyses emphasised the importance of the alpha-beta band in prediction effects, and discovered a putative role of beta-suppression in the encoding of precision of predictions. Our results highlight the effects of top-down information in the processing of affective stimuli and the importance of collecting behavioural data to characterize this relationship.

